# Host genotype and colonist arrival order jointly govern plant microbiome composition and function

**DOI:** 10.1101/2020.02.28.970582

**Authors:** Devin R. Leopold, Posy E. Busby

**Affiliations:** Department of Botany and Plant Pathology, Oregon State University, Corvallis, OR, 97331, USA

**Keywords:** community assembly, endophyte, foliar fungi, historical contingency, *Populus trichocarpa*, priority effects, plant microbiome

## Abstract

The composition of host-associated microbiomes can have important consequences for host health and fitness [1–3]. Yet, we still lack understanding of many fundamental processes that determine microbiome composition [4,5]. There is mounting evidence that historical contingency during microbiome assembly may overshadow more deterministic processes, such as the selective filters imposed by host traits [6–8]. More specifically, species arrival order has been frequently shown to affect microbiome composition [9–12], a phenomenon known as priority effects [13–15]. However, it is less clear whether priority effects during microbiome assembly are consequential for the host [16], or whether intraspecific variation in host traits can alter the trajectory of microbiome assembly under priority effects. In a greenhouse inoculation experiment using the black cottonwood (*Populus trichocarpa*) foliar microbiome, we manipulated host genotype and the colonization order of common foliar fungi. We quantified microbiome assembly outcomes using fungal marker-gene sequencing and measured susceptibility of the colonized host to a leaf rust pathogen, *Melampsora* × *columbiana*. We found that the effect of species arrival order on microbiome composition, and subsequent disease susceptibility, depended on the host genotype. Additionally, we found that microbiome assembly history can affect host disease susceptibility independent of microbiome composition at the time of pathogen exposure, suggesting that the interactive effects of species arrival order and host genotype can decouple community composition and function. Overall, these results highlight the importance of a key process underlying stochasticity in microbiome assembly while also revealing which hosts are most likely to experience these effects.

## Results and discussion

In this study, we sought to determine whether intraspecific variation in the model tree species, *Populus trichocarpa*, and the inoculation order of non-pathogenic foliar fungi interactively determine foliar microbiome composition and plant disease susceptibility (Figure 1). Specifically, we conducted a greenhouse experiment using 12 *P. trichocarpa* genotypes and a synthetic community of 8 common foliar fungi, which were isolated from field collected *P. trichocarpa* leaves. The plant genotypes can be broadly classified into two ecotypes, eastern and western, originating from either side of the Cascade Range that bisects the tree’s native range in Northwestern North America (Figure S1). The mountains create a sharp transition from a wet and mild environment in the west, to a dry continental environment in the east. The *P. trichocarpa* ecotypes associated with these regions vary in leaf morphology [17], foliar disease susceptibility [18–20], and are exposed to distinct microbial species pools [21]. Because these characteristics define the physical environment of the foliar microbiome (i.e., the leaf) [22,23], and suggest differences in innate immune responses to microbial colonization [24,25], we expected to observe the largest differences in host effects on foliar microbiome assembly when comparing eastern and western ecotypes.

**Figure 1:**
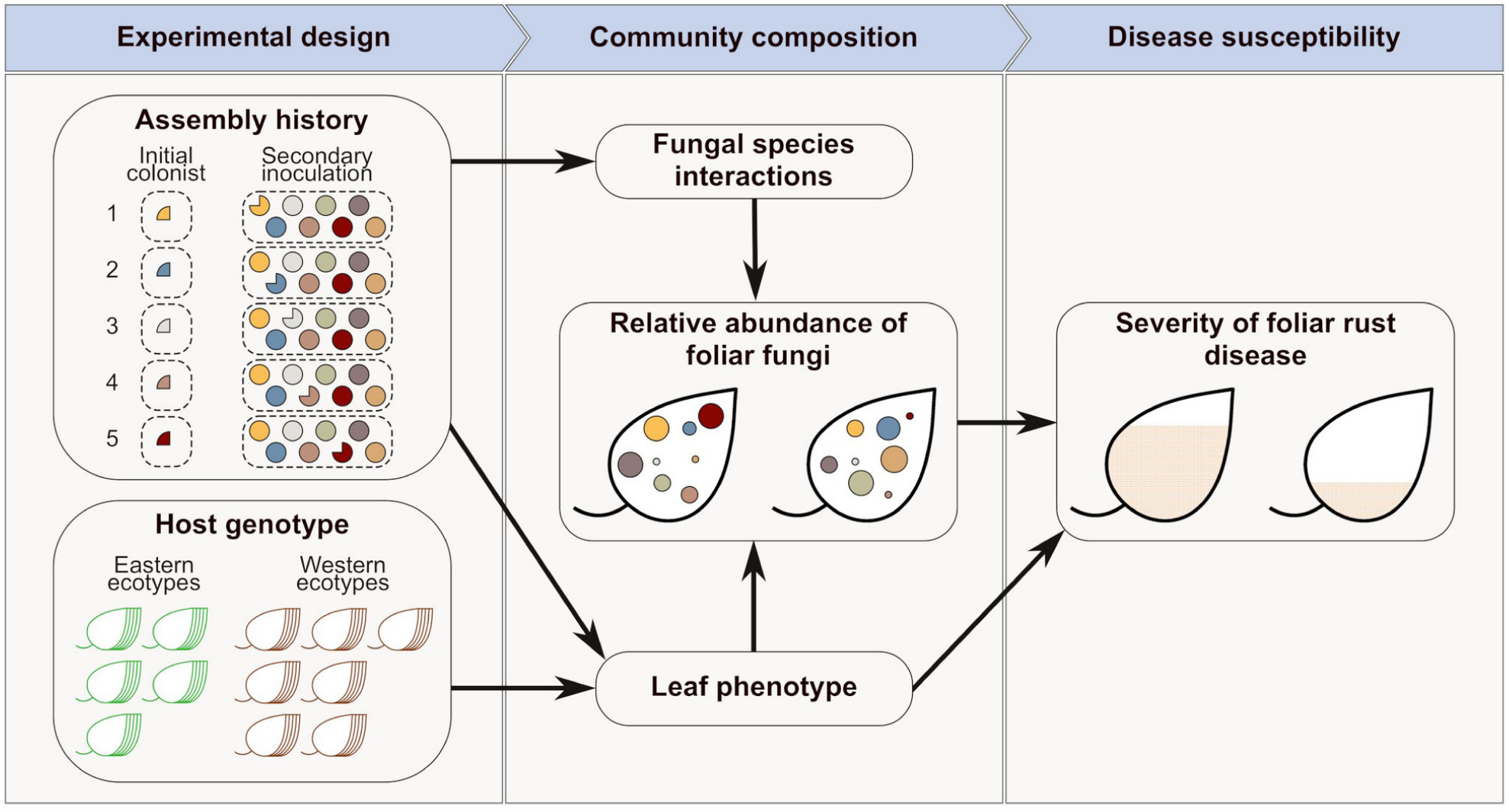
Experimental design and conceptual model. We factorially manipulated fungal community assembly history and host genotype. Assembly history treatments varied the identity of the initial colonist, for which one third of the total inoculum was applied preemptively. The 12 host genotypes included ecotypes originating from east and west of the Cascade Divide in Northwest North America. The identity of the initial colonist was hypothesized to affect microbiome composition by changing the outcomes of fungal species interactions during community assembly or by modifying the leaf phenotype. Host genotype was hypothesized to affect microbiome composition directly via host traits, such as leaf morphology or chemistry. We also hypothesized that both host genotype and assembly history could influence foliar rust disease severity via changes in microbiome composition or the host phenotype (e.g., modulation of plant defenses).

### Fungal species composition in the P. trichocarpa foliar microbiome depends on both species arrival order and host genotype

We first sought to test the hypothesis that the effect of species arrival order on the relative abundance of foliar fungi would vary among *P. trichocarpa* genotypes and ecotypes. We created 5 immigration history treatments in which one member of our 8-species synthetic community was allowed to preemptively colonize leaves, followed by inoculation with the full community 2 weeks later. We assessed fungal community assembly outcomes using marker-gene sequencing and modeled variation in species relative abundances using joint-species distribution models [26], accounting for taxon-specific sequencing biases using empirical estimates derived from mock-community data (Figures S2).

We found that the relative abundances of foliar fungi varied across the experimental treatments, with some species responding more strongly to species inoculation order and others responding more strongly to host genotype (Figures 2A and 2B). The effect of species arrival order was not limited to taxa that served as initial colonists, indicating that the identity of the initial colonist can result in complex indirect effects on the trajectory of community assembly. However, when the relative abundances of individual species were analyzed independently (Figure 2B) we did not find strong evidence that the effect of species arrival order varied among host genotypes; an effect that only became evident for the community as a whole (see below).

**Figure 2:**
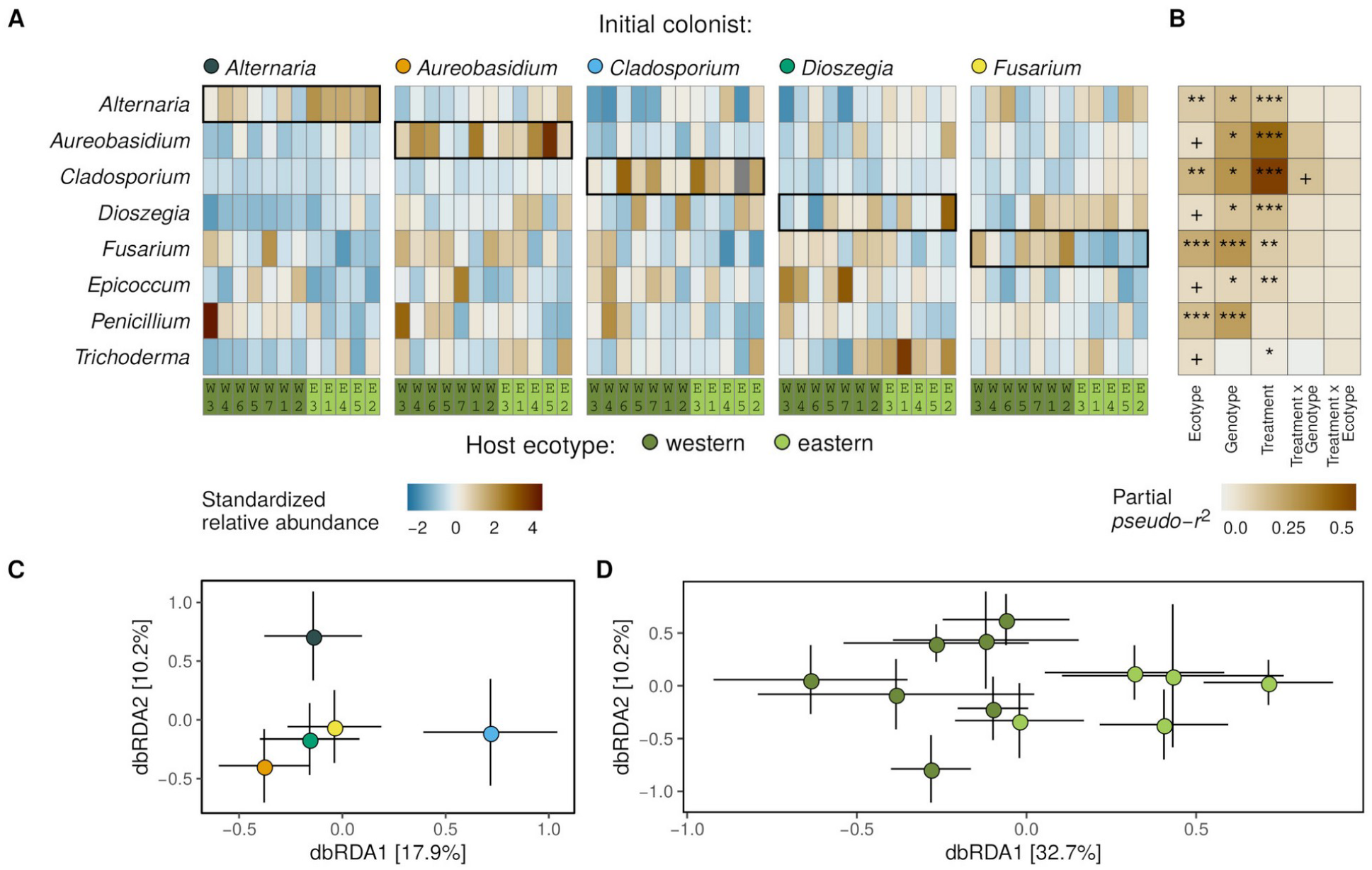
Fungal species composition in the foliar microbiome depends on both species arrival order and host genotype. (A) Heatmap showing variation in fungal relative abundance for each species (rows) and plant genotype (columns). Panels show the 5 immigration history treatments, labeled with the identity of the initial colonist. Highlighted cells show the relative abundance of the initial colonist in each immigration history treatment. Columns representing individual host genotypes in each panel are organized according to their position along the first axis of the constrained ordination in D. (B) Results of univariate models of individual species relative abundances, testing the joint effects of species arrival order treatments, host genotype, and host ecotype (For associated community-level results see Table S1). Colors indicate the partial *pseudo*-*r*^2^ for each model term, calculated following the method of Nagelkerke [58]. Symbols indicate the significance (permutation tests) of individual species’ contributions to the multivariate (community-level) response, where ^+^*P* < 0.1, ^*^*P* < 0.05, ^**^*P* < 0.01, and ^***^*P* < 0.001. (C) Distance-based redundancy analysis, showing variation in fungal community dissimilarity (Jensen-Shannon distance) among initial colonist treatments after conditioning on the effect of host genotype. Colored points show the group means (± sd) for initial colonist treatments on the first 2 constrained axes. (D) Distance-based redundancy analysis, showing variation in fungal community dissimilarity (Jensen-Shannon distance) among host genotypes after conditioning on the effect of the initial colonist treatments. Colored points show the group means (± sd) for each plant genotype on the first 2 constrained axes and color indicate host ecotype.

At the community-level, we found that fungal species composition depended on species arrival order (Wald-*χ*^2^_(4)_ = 21.8, *P* < 0.001; Figure 2C), and that the effect of arrival order varied across plant genotypes (Wald-*χ*^2^_(44)_ = 23.5, *P* = 0.04; Table S1). The direct effect of plant genotype on fungal species composition (Wald-*χ*^2^_(11)_ = 20.2, *P* < 0.001) was partially explained by host ecotype (Wald-*χ*^2^_(1)_ = 12.4, *P* < 0.001; Figure 2D). However, contrary to our expectations, we did not find evidence to support the hypothesis that the effect of species arrival order on fungal community composition is systematically influenced by host ecotype (Wald-*χ*^2^_(4)_ = 6.16, *P* = 0.31; Table S1). This suggests that the plant traits that interact with species arrival order to affect community assembly are not the physical leaf traits that broadly segregate across the wet and dry regions sampled. It is possible that other traits with greater variance among individual *P. trichocarpa* genotypes, such as those involved in microbial recognition and immune response signaling [27–29], are responsible for the genotype-by-arrival order interaction we observed. However, genome wide association studies across many more plant genotypes are needed to test this hypothesis [30].

### The advantage of preemptive colonization depends on species identity and host genotype, but not host ecotype

Variation in the identity of the initial colonist affected the relative abundances of all but one member of the synthetic community of foliar fungi (Figures 2A and B), yet, only 3 of the 5 early arriving species consistently experienced a benefit from preemptive colonization (Figure 3). This could be the result of intrinsic fitness differences among the initial colonists, or differences in niche overlap with the other community members, affecting the benefit of early arrival [31,32]. For example, *Alternaria* had the greatest relative abundance across all treatments and, although it did benefit from early arrival, the magnitude of priority effects for this species was relatively low. This could suggest that *Alternaria* is competitively dominant within the foliar microbiome, limiting the possibility of niche preemption by other species. However, we cannot exclude the possibility that this species occupies a distinct niche within the leaf, particularly given that the genus *Alternaria* is more distantly related to the other isolates that benefited from early arrival, *Cladosporium* and *Aureobasidium*, then they are to one another [33,34]. Niche differences, such as specialization on the leaf surface vs. interstitial space would also reduce the relative importance of arrival order and could explain why species that did not benefit from early arrival often varied in relative abundance in response to the identity of other initial colonists.

**Figure 3:**
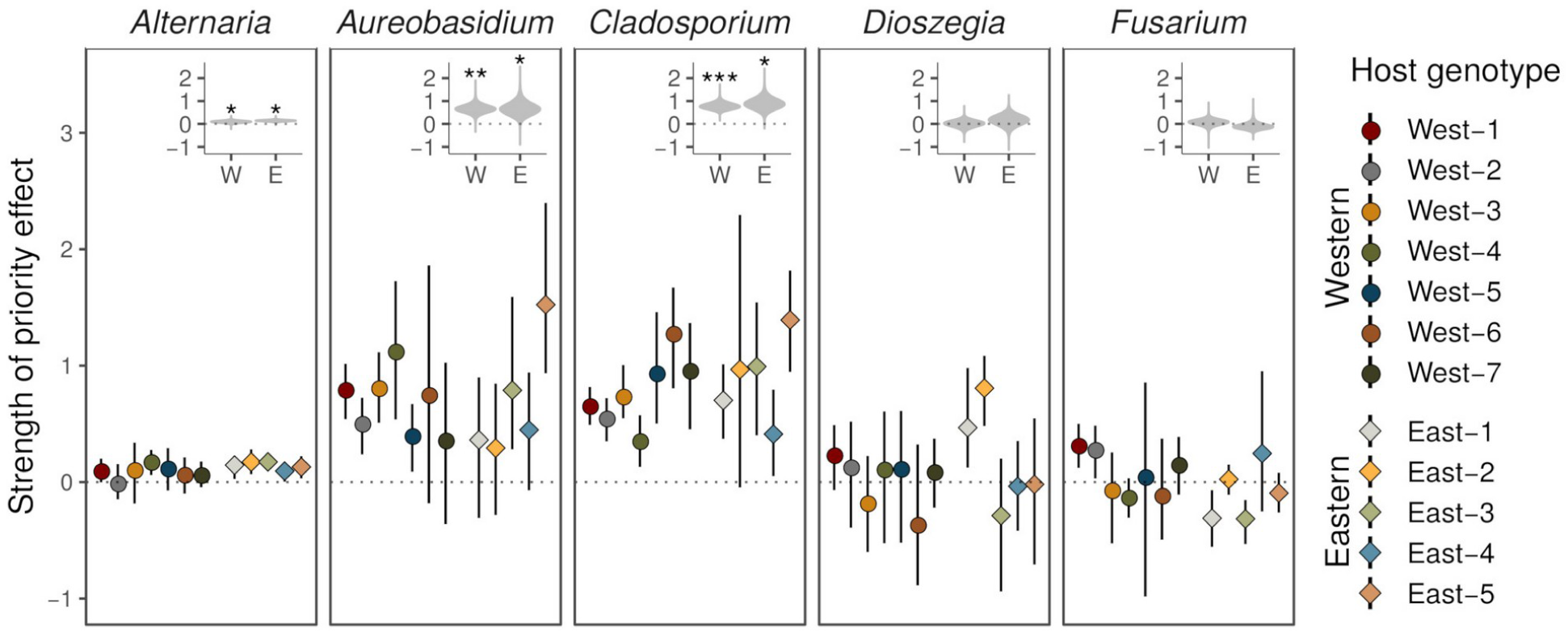
The advantage of preemptive colonization in the foliar microbiome depends on species identity and host genotype, but not host ecotype. The strength of priority effects for each species on each genotype was calculated as the log ratio of the relative abundance when arriving first vs. arriving concurrent with the community. Points show the estimated priority effect for each fungal species (panels) on each plant genotype (point colors) and error bars indicate 95% bootstrapped confidence intervals. Insets show bootstrap distributions of the mean strength of priority effects for the eastern and western *P. trichocarpa* ecotypes. Significant mean priority effects, based on bootstrapped one-sample t-tests, are indicated, where ^*^*P* < 0.05, ^**^*P* < 0.01, and ^***^*P* < 0.01.

Because the synthetic communities used in this experiment all included the same 8 fungal species, yet natural plant microbiomes can host hundreds species [35,36], it is likely that our experimental design represents a conservative test for the strength of priority effects. Priority effects are more likely to be strong when the pool of potential colonists is larger because greater niche overlap and niche modification are both more likely [37]. In addition, our fungal isolates were collected from multiple locations, which were not the same *P. trichocarpa* stands where the plant genotypes originated (Figure S1). If local adaptation to host populations plays a role in plant microbiome assembly [38], priority effects could be stronger within pools of locally adapted symbionts [39,40]. Finally, our immigration history treatments involved applying just one third of the total inoculum of one species initially, with the remainder applied concurrent with the other community members. This was done to ensure that any observed effects were the result of preemptive colonization and not differences in total quantity of inoculum or absence of the initial colonist from the second inoculation. As a result, our assembly history perturbations were relatively subtle compared to the potential stochasticity that can result from dispersal processes occurring in natural microbial communities [41].

### Foliar microbiome assembly history affects rust disease severity in susceptible plant genotypes

To test the hypothesis that foliar disease severity depends on the joint effects of host genotype and species arrival order during foliar microbiome assembly, we inoculated plants with the leaf rust pathogen, *Melampsora* × *columbiana*, 2 weeks after establishing foliar microbiomes under our immigration history treatments. We then used image analysis to quantify disease severity as the proportion of leaf area that developed chlorotic lesions and used beta-regression models to assess variation across treatments (Figure 4 & Table S2). As expected, the primary source of variation in rust disease severity was host ecotype, likely reflecting the presence of major-gene resistance to the rust pathogen in most, but not all, western genotypes (Figures 4 & S3).

**Figure 4:**
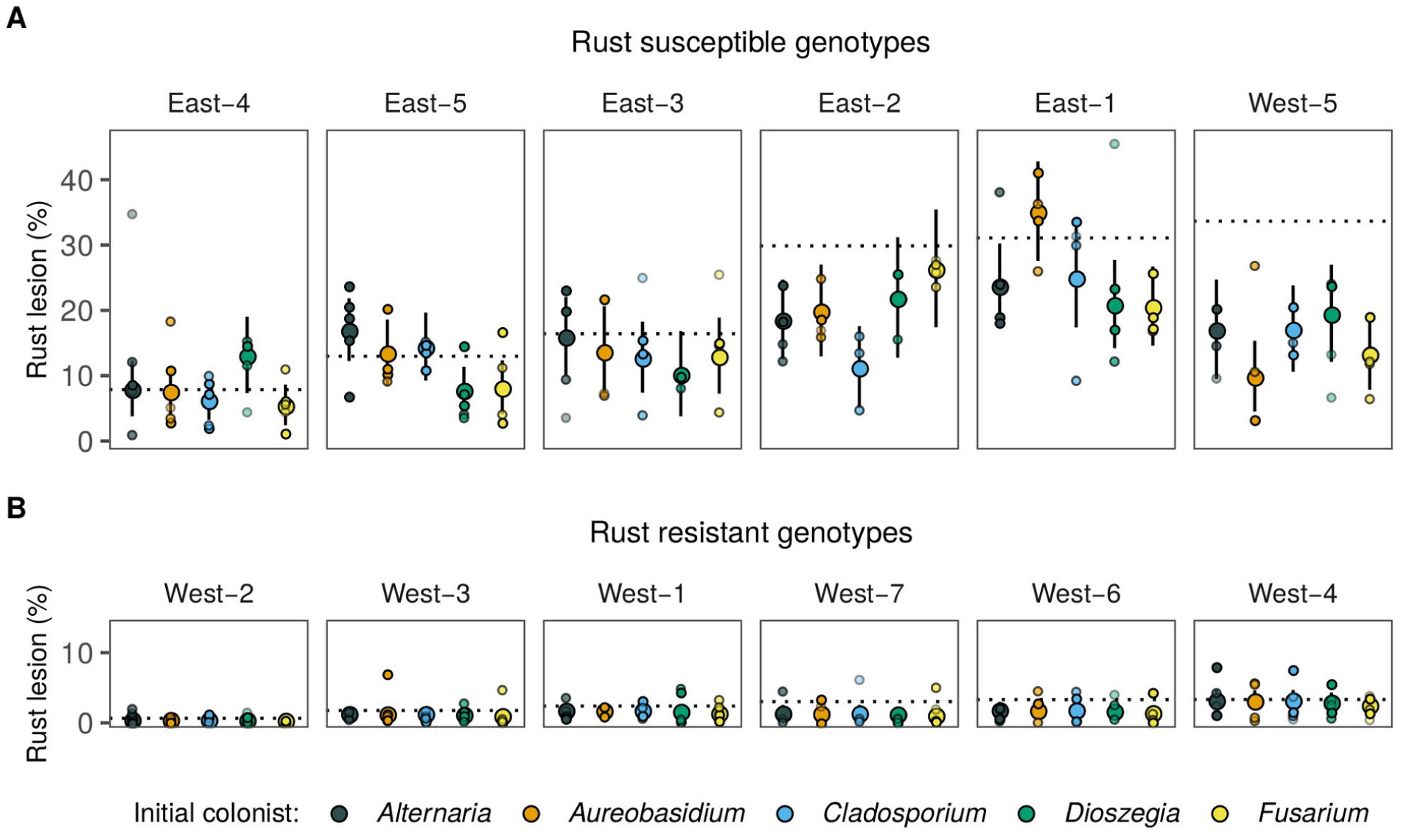
Foliar microbiome assembly history affects rust disease severity in susceptible plant genotypes. Large points indicate estimated marginal means (95% CIs) from beta-regression models of rust disease severity in (A) susceptible and (B) resistant *P. trichocarpa* genotypes. Small points show individual observations for each species arrival order treatment (colors) on each host genotype (panels). Transparency of small points indicates the weight used (number of individual leaves measured per plant) in the beta-regression models. Horizontal dotted lines indicate the mean rust severity in uninoculated plants (Figure S3). The results of the beta-regression models are presented in Table S2. The relationship between rust severity and the relative abundance of foliar fungi is presented in Figure S4.

We found that the effect of fungal species arrival order on rust disease severity depended on whether the host genotype was susceptible or resistant to the rust pathogen. Specifically, for susceptible, primarily eastern genotypes, disease severity depended on the interaction between fungal species arrival order and host genotype (LRT-*χ*^2^ _(20)_ = 33.6, *P* = 0.03; Figure 4A). However, while there was variation in the overall level of rust disease observed among the rust resistant, western genotypes (LRT-*χ*^2^_(5)_ = 33.5, *P* < 0.001), the arrival order of foliar fungi did not affect rust disease development (LRT-*χ*^2^ _(20)_ = 1.74, *P* = 0.78; Figure 4B).

Although we found that microbiome assembly history affected subsequent disease severity in rust-susceptible plant genotypes, we did not find evidence that this was the result of changes in the relative abundance of individual fungi or overall shifts in foliar microbiome composition (Figure S4). One explanation for this could be that disease modification by early arriving species occurred primarily through interactions with the plant’s immune system [42,43], decoupling the outcome of microbiome community assembly from disease development (Figure 1). Although some foliar fungi can have large effects on *P. trichocarpa* foliar rust severity [44], we did not pre-screen fungi to identify species that were associated with disease modification. Including such species in our synthetic communities may have yielded stronger links between fungal community composition and rust disease. In addition, we did not systematically sample co-occurring plants and fungi. Local adaptation between microbial symbionts and plants could result in non-additive contributions of plant and fungal genotypes to host disease susceptibility, analogous to the genotype-by-genotype interactions that can arise from host-pathogen co-evolutionary dynamics [45].

Standard approaches for marker-gene sequencing of microbiome communities only provide data on the relative abundances of taxa within each sample, i.e., compositional data [46,47]. It is possible that total fungal abundance in the foliar microbiome varies among plant genotypes due to variation in resource availability, leaf thickness, or other leaf traits [48]. Even the identity of the initial colonist could affect subsequent microbial abundance in leaves if host defenses are triggered or suppressed [49]. Variation in total fungal abundance could be consequential for disease modulation, particularly if direct pathogen antagonism by non-pathogenic microbes contributes to disease suppression. Variation in total microbial abundance could also have implications for priority effects during microbiome community assembly. Specifically, priority effects should be more important when space or resources limit the community size because niche preemption by early arriving species is more likely [50,51]. However, priority effects may be stronger in resource rich environments if the growth rate of early arriving species is increased [52,53], though greater local resource availability can reduce the strength of priority effects in some fungi by reducing the rate of substrate exploration [54]. Progress towards understanding the interactive effects species arrival order and host genotype on plant microbiome composition and function will require quantitative data on microbial abundance.

While manipulations of synthetic plant microbiomes is increasingly seen as a valuable experimental approach, most efforts have focused on bacteria [11,55,56]. However, fungi are often the most functionally prominent plant symbionts [57]. Our study has demonstrated the feasibility of controlled manipulations of fungal community assembly in the phyllosphere using a tractable synthetic community. Using this approach, we have shown that the impacts of fungal immigration history on microbiome composition and plant disease susceptibility depend on plant genotype. Future experiments investigating the sensitivity of fungal immigration history to a broader range of host and environmental variation will establish the foundational knowledge needed to realize the promise of selective manipulation of plant microbiomes for applications in agriculture or natural resource management.

## Acknowledgments

We are grateful to Bruce McCune, George Newcombe, Maggie Wagner, and Timothy Warren for feedback during the preparation of this manuscript. This research was supported by the US Department of Energy’s Office of Biological and Environmental Research (DE-SC0019435) and by the Oregon State University, Department of Botany and Plant Pathology.

## Author contributions

DRL and PEB jointly planned the study. DRL lead implementation, data collection, analysis, and manuscript preparation, with input and contributions from PEB at all stages.

## Star methods

### Resource Availability

#### Lead contact

Requests for resources of further information should be directed to the Lead Contact, Devin R. Leopold.

#### Materials availability

This study did not generate new reagents.

#### Data and code availability

The high-throughput sequence data generated in this study is available at the NCBI Sequence Read Archive (BioProject PRJNA605581). All code used to process data and generate the results presented in this study, as well as sample meta data, is available as a publicly archived git repository (DOI: 10.5281/zenodo.3872145).

### Experimental model and subject details

#### Populus trichocarpa

Plants were propagated in a glasshouse on the campus of Oregon State University using dormant hardwood cuttings collected in February 2018, from a common garden. Cuttings were trimmed to *ca.* 10 cm and planted into 550 ml conical Deepots (Stuewe & Sons, Inc.) using commercial potting mix (Sunshine Mix #4) supplemented with 115 ml gal^-1^ slow-release fertilizer (16-6-11 + micronutrients). Throughout the experiment, plants were watered from below using an automated sub-irrigation system to keep leaves dry and limit opportunities for colonization by non-target fungi [59].

#### Fungal cultures

We isolated foliar fungi from *P. trichocarpa* leaves using standard methods. Briefly, leaf punches were collected into sterile 1.5 ml centrifuge tubes and surface sterilized by sequentially vortexing for 1 minute in 10% bleach (0.825% NaClO), 70% EtOH, and then in 3 consecutive rises in sterile water. We then plated the leaf punches onto potato-dextrose agar (PDA), supplemented with penicillin (100 μg ml^-1^) and streptomycin (100 μg ml^-1^), and incubated the plates at room temperature for 1 month. Plates were checked daily for fungal growth and emerging hyphae were subcultured using single hyphal-tip isolation. Cultures were maintained during the experiment by serial transfer on PDA and then archived for long-term storage in sterile water at 4 °C.

### Method details

#### Overview and experimental design

We manipulated the species arrival order of foliar fungi colonizing the leaves of 12 genotypes of the model tree species, *P. trichocarpa*, in a controlled greenhouse experiment. The plant genotypes originated from populations located both east (5) and west (7) of the Cascade Divide in the northwestern United States (Figure S1). In the greenhouse, plants were inoculated with a synthetic community of 8 fungal species (Table S3), all isolated from field collected *P. trichocarpa* leaves (Figure S1). Inoculation consisted of two phases: colonization by 1 of 5 randomly selected initial colonists, followed 2 weeks later by inoculation with the full synthetic fungal community. The 5 inoculation treatments, and an additional non-inoculated control treatment, were applied to 5 replicates of 12 clonally propagated *P. trichocarpa* genotypes, resulting in a total of 360 plants. We assessed fungal community assembly outcomes after one month using Illumina sequencing of the nuclear ribosomal, internal transcribed spacer (ITS) region. We then inoculated plants with the foliar rust pathogen *Melampsora × columbiana* and assessed disease severity to quantify functional consequences of priority effects in the *P. trichocarpa* foliar microbiome.

#### Synthetic fungal community

We selected representative morphotypes from the initial fungal culture collection (see *Experimental model and subject details*, above) and used Sanger sequencing of the full ITS region for taxonomic identification. We then selected representative isolates of 8 fungal species (Table S3) to use in our synthetic community of *P. trichocarpa* foliar fungi, which included: *Alternaria alternata, Aureobasidium pullulans, Cladospoium* sp., *Dioszegia butyracea, Epicoccum nigrum, Fusarium* sp., *Penicillium bialowiezense*, and *Trichoderma trixiae*. The isolates we selected have been previously identified as common members of the *P. trichocarpa* foliar microbiome [21,44]. Because definitive species-level identifications from ITS sequence data is not always possible, the genus-level taxonomic assignments are used to refer to the individual isolates throughout this study. Sanger sequences of the full ITS region for the 8 focal species are archived in the NCBI GenBank (accession numbers MT035960-7).

#### Fungal inoculation

We collected fungal inoculum by flooding 2-week old cultures with an aqueous solution of 0.1% Tween 20 and gently scraping the plate surface with a sterile spatula to dislodge conidia, or, in the cases of the yeast-like *Aureobasidium pullans* and the basideomycetous yeast, *Dioszegia* sp., individual cells. We quantified the density of the initial suspension using a hemocytometer and diluted as required using additional sterile 0.1% Tween 20. At the first inoculation time point, we prepared inoculation solutions for each of the 5 initial colonists at a density of 3.3 x 10^4^ cells ml^-1^. At the second time point, 2 weeks later, inoculation solutions for each treatment were prepared containing each of the 7 other species at a density of 1.0 x 10^5^ cells ml^-1^ and the initial colonist at a density of 6.7 x 10^4^ cells ml^-1^. As a result, the total amount of inoculum for each species in each treatment was held constant and only the identity of the initial colonist varied. We chose to apply the majority of the inoculum for the initial colonist at the second time point to ensure that differences in assembly outcomes were due to early arrival, and not the absence of the initial colonist from the inoculation solution during colonization by the full community.

Inoculation of plants was achieved by saturating both the top and bottom leaf surfaces with a sterile, fine-mist spray bottle. For each tray of 12 plants, we applied 75 ml of the inoculum solution and then immediately covered the tray with a temporary humidity tent to maintain leaf surface moisture for 48 hours. Control plants were mock-inoculated using sterile water at both time points. Because *P. trichocarpa* grows rapidly and the experiment was conducted over a period of 6 weeks, we used a wire twist tie to mark the youngest fully expanded leaf at the time of the first inoculation.

#### Leaf sampling for fungal DNA

From each plant, we collected 4 leaf punches (*ca*. 0.25 cm^2^) from each of 3 leaves (12 total) directly into a sterile 1.5 ml centrifuge tube using a standard hole punch, which was cleaned with ethanol between each plant. Leaf disks were placed on ice immediately after collection and processed to remove surface contaminants the same day they were collected. To remove surface contaminants, the leaf disks were agitated for 1 minute in 1 ml of molecular grade water with 0.1% Tween 20, using a Geno/Grinder® 2010, set at 500 strokes per minute. The cleaning solution was remove from the tubes using a sterile pipette and the leaf disks were rinsed by repeating the process 3 times with molecular grade sterile water. The cleaned leaf disks were then frozen at −20 °C and stored until we extracted total genomic DNA using the 96 Well Synergy™ Plant DNA Extraction Kit (OPS Diagnostics). The four corners of each extraction plate were processed without leaf disks to serve as negative controls during library preparation.

#### Library preparation for ITS metabarcoding

We prepared libraries for Illumina MiSeq metabarcoding, targeting the ITS1 region using a dual-indexed, two-stage PCR approach [60]. In the first PCR, we used modified versions the fungal specific forward gene primer ITS1F [61] and, for the reverse gene primer, the reverse-complement of gITS7 [62]. Each gene primer was fused with Illumina sequencing primers, separated by a 3-6 bp, degenerate, length-heterogeneity spacer [63]. Stage-one PCR was carried out in 25 µl reactions with MyFi Master Mix (Bioline), 2 µl template DNA, 0.5 µM of each primer and a thermocycling program consisting of an initial denaturing and enzyme activation cycle at 95 °C (3 min.), 32 cycles of 95 °C (30 sec.), 50 °C (30 sec.), and 72 °C (30 sec.), followed by a final elongation cycle at 72 °C (5 min). Temperature ramp rates were limited to 1 °C [64]. For each 96-well plate, 2 extraction blanks were included to detect contaminants possibly introduced during DNA extraction and 2 PCR blanks, with no template DNA, were included to detect contamination introduced during library preparation. For stage-two PCR, we used primers that targeted the Illumina sequencing adapters added during stage-one, adding the P5 and P7 Illumina adapters and 8-mer multiplexing barcodes [65]. Multiplexing barcodes were selected to have a minimum Hamming distance of 4 and balanced base diversity [66]. Stage-two PCR was carried out in 20 µl reactions with MyFi Master Mix, 1 µl of the stage-one PCR product as template, 0.5 µM primers, and a thermocycler program consisting of an initial cycle of 95 °C (1 min.), 8 cycles of 95 °C (20 sec.), 55 °C (20 sec.), and 72 °C (30 sec.), followed by a final elongation cycle at 72 °C (5 min). Final PCR products were cleaned and normalized using Just-a-Plate, 96-well normalization and purification plates (Charm Biotech), then pooled and sent to the Oregon State University, Center for Genome Research and Biocomputing, for 250-bp, paired-end sequencing on the Illumina MiSeq platform.

#### Fungal community bioinformatics

Sequences were demultiplexed with Pheniqs v2.0.4 [67], using a phred-adjusted maximum likelihood confidence threshold of 0.995. Gene primers and length heterogeneity spacers were removed from the 5’ ends of the paired reads using cutadapt v1.18 [68], discarding reads with gene primers not found in the expected positions. The 3’ ends were trimmed to remove “read-through” adapter contamination using SeqPurge [69]. Following trimming, reads were filtered at a maximum expected error rate of 2 bp, denoised, and filtered of putative chimeras using DADA2 [70], with the expected amplicons for the 8 members of the synthetic community included as denoising priors to increase sensitivity. Denoised amplicons were then further collapsed to operational taxonomic units (OTUs) at 99% similarity using agglomerative, single-linkage clustering with the R-package DECIPHER [71] used to calculate pairwise dissimilarity. Host contamination was removed by filtering the resulting OTUs against the *P. trichocarpa* v3.0 genome [72] using Bowtie2 [73]. Other likely contaminants were filtered by removing OTUs whose prevalence or mean proportional abundance when present was greater in negative controls. Samples with less than 4000 sequences remaining after all filtering steps were removed from subsequent analyses. The final OTU-by-sample matrix and the sample metadata were combined in a single phyloseq object [74] for downstream manipulation.

#### Estimating sequencing bias

To quantify and correct for species-specific sequencing biases introduced during Illumina library preparation and sequencing, we constructed mock communities with known abundances of fungal DNA that were sequenced along with our experimental samples. To assemble the mock communities, we first extracted genomic DNA from pure cultures of each of the 8 members of the synthetic community and the rust pathogen using the *Quick*-DNA Fungal/Bacterial Miniprep Kit (Zymo Research). We then used Sanger sequencing and Primer 3 [75] to design custom qPCR primers to quantify the concentrations of our pure culture extracts. We targeted the TEF1-α gene for qPCR quantification because it is a single-copy gene, unlike the ITS region, which can vary in copy number among fungi over several orders of magnitude [76], likely representing a large source of bias in fungal ITS metabarcoding. Absolute qPCR quantification was carried out using PowerUP SYBR Master Mix (Applied Biosystems) and the 7500 Fast Real-Time PCR System (Applied Biosystems). We then used the qPCR estimated concentrations of the single culture extracts to combine genomic DNA for each species in 10 different mock communities, each containing all 9 fungal species. For 1 mock community, all species were present in equal concentrations (5 x 10^4^ copies µl^-1^). For the remaining 9 mock communities, species were added at either, 1 x 10^4^ copies µl^-1^, 5 x 10^4^ copies µl^-1^, or 2.5 x 10^5^ copies µl^-1^. Each species was present at each concentration in 3 mock communities and all 9 mock communities had 3 species at each concentration. The resulting mock community samples were prepared for Illumina sequencing in parallel with the experimental samples following the protocol outlined above.

Sequencing read counts and the known proportional abundances of each species in each mock community were then used to estimate species-specific biases following the method of [77]. We found significant bias among the members of our mock communities (Figure S2), with *M.* x *columbiana*, having the greatest positive bias, being over represented relative to the most negatively biased species, *Penicillium* sp. by a factor of 40. Among the members of our synthetic community of foliar fungi, the most positively biased species was *Fusarium* sp., which was over represented relative to *Penicillium* sp. by a factor of 13. Application of the bias estimates from mock community data to account for sequencing bias in experimental samples is described in the Quantification and statistical analysis section below.

#### Rust disease severity

Immediately following leaf sampling for marker-gene sequencing, we inoculated plants with a rust pathogen. Asexual urediniospores were collected from a single-spore strain of *M. × columbiana* being maintained on *P. trichocarpa* plants in an isolated growth chamber. We applied urediniospores at a concentration of 1.0 x 10^4^ ml^-1^, following the same procedure described above. After 2 weeks, we harvested 3 leaves from each plant and photographed the abaxial surface from a fixed distance using a light box. We then used image analysis, with Fiji [78], to quantify rust disease severity. First, we isolated leaves from the image background and quantified total leaf area using color thresholding. We then used supervised image segmentation [79] to quantify the proportion each leaf occupied by either chlorotic lesions or rust uredinia, to related measures of rust disease severity.

### Quantification and statistical analysis

To assess whether the community assembly of *P. trichocarpa* foliar fungi was influenced by the 5 colonization order treatments, 12 host genotypes, and their interaction, we fit a multivariate, negative-binomial glm, using the R-package mvabund [80]. We accounted for 2 sources of unequal sampling effort, variable sampling depth and the species-specific sequencing biases, by including an offset term (*effort*) for each species *i* in each sample *j*, in the form: *effort*_*ij*_ = log(*bias*_*i*_ x *depth*_*j*_), where *bias*_*i*_ is the sequencing bias correction factor for species *i*, estimated from our mock communities (see above, *Estimating Sequencing Bias*), and *depth*_*j*_ is the total sum of all species in sample *j*, after dividing each by their species-specific sequencing bias correction factor. We used Wald tests to asses the significance of model terms, accounting for non-independence of species relative abundances by estimating a species correlation matrix [81] and by assessing significance using permutation tests, while maintaining species correlations, to calculate p-values [82]. To determine whether host ecotype explained the effects of host genotype, we fit a second multivariate glm with plant genotype re-coded as a binary factor (eastern or western).

We used two approaches to determine the relative contributions of individual species to the overall variation in community composition in response to our experimental treatments. First, we extracted univariate test statistics from our multivariate glms, using permutation tests and step-down resampling to control the family-wise error rates. To quantify univariate effect sizes, we fit individual negative-binomial glms for each species with the R-package MASS [83], and estimated the partial-*R*^2^ for each model term, following the method of Nagelkerke [58].

As an alternative approach to assessing the impact of the experimental treatments on individual species, we quantified the benefit of preemptive colonization (i.e., priority effects) for each of the 5 initial colonists. For each initial colonist on each host genotype, we defined the strength of priority effects as the log-ratio of the mean proportional abundance when arriving first to the mean proportional abundance when arriving concurrent with the community, following [84]. This calculation results in positive values if the relative abundance was greater when the species arrived before the community. We calculated point estimates for the strength of the priority effect for each species on each host genotype, using bootstrapping to estimate confidence intervals, randomly sampling with replacement 10,000 times for each treatment combination and recalculating the strength of the priority effect for each iteration. Bias-corrected and accelerated confidence intervals [85] were estimated from the resulting distributions. To test whether each species experienced significant priority effects on eastern or western ecotypes, we used bootstrapped one-sample t-tests. To ensure that the bootstrapping process resembled the data generation process, we used a two-stage resampling procedure, beginning with the 10,000 bootstrapped point-estimates of priority effects describe above. For each set of bootstrapped point estimates, we drew 100 bootstrap samples for each fungal species and host ecotype combination and calculated the t-statistic under the null hypothesis of no difference in relative abundance due to arrival timing. The bootstrapped, one-tailed p-values were then calculated as the proportion of times the bootstrapped t-statistics were greater than the values from the observed data.

We then tested whether our experimental treatments affected the severity of leaf rust disease, measured as the proportion of leaf surface occupied by chlorotic lesions or rust uredinia, averaged over three leaves per plant. We separated highly resistant and susceptible genotypes for this analysis because our disease severity data indicated 2 qualitatively distinct groups, one of which likely posses major-gene resistance to the rust pathogen [29,86], precluding variation in disease severity (Figure S3). For each group, we then fit generalized linear models with a beta distribution (i.e., beta-regression), using the R-package betareg [87]. To account for the presence of zeros in the rust uredinia data, a transformation was applied, following [88], (*y* * (*n* - 1) + 0.5) / *n*, where *y* is the proportion of leaf with rust uredinia and *n* is the total number of plants sampled. In addition, because some individual leaves were lost during handling, we accounted variation in measurement precision by using the number of leaves sampled per plant as proportionality weights in the models. To determine whether host genotype and fungal species arrival order interactively determined disease severity, we compared models with and without an interaction term using likelihood-ratio tests, implemented with the R-package lmtest [89]. We then tested the marginal direct effects of host genotype and species arrival order on disease severity. Analyses of both measures of rust disease severity, chlorotic lesions and uredinia, yielded identical conclusions, so we present only the former.

We then tested whether variation in disease severity within each susceptible plant genotype could be explained by variation in the relative abundance of individual foliar fungi, or community-level variation associated with the species arrival order treatments. First, we fit a new beta-regression model to the rust severity data with only host genotype as a predictor. We then extracted the standardized residuals to use as a new response variable capturing variation in disease severity after accounting for differences among individual plant genotypes. To assess whether residual rust severity was related to the relative abundance of individual foliar fungi, we then conducted non-parametric correlation tests (Kendall’s tau) and examined scatter plots, using log-odds transformed proportional fungal abundance. To assess whether residual rust severity was associated with the species arrival order treatment effects on overall fungal community composition, we first conducted a distance based redundancy analysis of fungal community dissimilarity, using Jensen-Shannon Distance, and conditioned on host genotype effects, to identify orthogonal axes of variation associated species arrival order. We then used non-parametric correlation tests and examined scatter plots, as above.

All analyses were conducted using R v3.6.2 [83].

## References

1. Vandenkoornhuyse, P., Quaiser, A., Duhamel, M., Le Van, A., and Dufresne, A. (2015). The importance of the microbiome of the plant holobiont. New Phytol. 206, 1196–1206.

2. Laforest-Lapointe, I., Paquette, A., Messier, C., and Kembel, S.W. (2017). Leaf bacterial diversity mediates plant diversity and ecosystem function relationships. Nature 546, 145–147.

3. Cho, I., and Blaser, M.J. (2012). The human microbiome: at the interface of health and disease. Nat. Rev. Genet. 13, 260–270.

4. Tripathi, A., Marotz, C., Gonzalez, A., Vázquez-Baeza, Y., Song, S.J., Bouslimani, A., McDonald, D., Zhu, Q., Sanders, J.G., Smarr, L., et al. (2018). Are microbiome studies ready for hypothesis-driven research? Curr. Opin. Microbiol. 44, 61–69.

5. Lajoie, G., and Kembel, S.W. (2019). Making the most of trait-based approaches for microbial ecology. Trends Microbiol. 27, 814–823.

6. Ricci, F., Rossetto Marcelino, V., Blackall, L.L., Kühl, M., Medina, M., and Verbruggen, H. (2019). Beneath the surface: community assembly and functions of the coral skeleton microbiome. Microbiome 7, 1–10.

7. Sprockett, D., Fukami, T., and Relman, D.A. (2018). Role of priority effects in the early-life assembly of the gut microbiota. Nat. Rev. Gastroenterol. Hepatol. 15, 197–205.

8. Moccia, K.M., and Lebeis, S.L. (2019). Microbial ecology: how to fight the establishment. Curr. Biol. 29, R1320–R1323.

9. Kennedy, P.G., Peay, K.G., and Bruns, T.D. (2009). Root tip competition among ectomycorrhizal fungi: are priority effects a rule or an exception? Ecology 90, 2098–2107.

10. Werner, G.D.A., and Kiers, E.T. (2015). Order of arrival structures arbuscular mycorrhizal colonization of plants. New Phytol. 205, 1515–1524.

11. Carlström, C.I., Field, C.M., Bortfeld-Miller, M., Müller, B., Sunagawa, S., and Vorholt, J.A. (2019). Synthetic microbiota reveal priority effects and keystone strains in the *Arabidopsis* phyllosphere. Nat. Ecol. Evol. 3, 1445–1454.

12. Martínez, I., Maldonado-Gomez, M.X., Gomes-Neto, J.C., Kittana, H., Ding, H., Schmaltz, R., Joglekar, P., Cardona, R.J., Marsteller, N.L., Kembel, S.W., et al. (2018). Experimental evaluation of the importance of colonization history in early-life gut microbiota assembly. Elife 7, 1–26.

13. Palmgren, A. (1926). Chance as an element in plant geography. In Proceedings of the International Congress of Plant Sciences, B. Duggar, ed. (Ithica, New York), pp. 591–602.

14. Sutherland, J.P. (1974). Multiple stable points in natural communities. Am. Nat. 108, 859–873.

15. Drake, J. (1991). Community-assembly mechanics and the structure of an experimental species ensemble. Am. Nat. 137, 1–26.

16. Litvak, Y., and Bäumler, A.J. (2019). The founder hypothesis: a basis for microbiota resistance, diversity in taxa carriage, and colonization resistance against pathogens. PLoS Pathog. 15, 1–6.

17. Dunlap, J.M., and Stettler, R.F. (2001). Variation in leaf epidermal and stomatal traits of *Populus trichocarpa* from two transects across the Washington Cascades. Can. J. Bot. 79, 528–536.

18. Dunlap, J.M., and Stettler, R.F. (1996). Genetic variation and productivity of *Populus trichocarpa* and its hybrids. IX. Phenology and Melampsora rust incidence of native black cottonwood clones from four river valleys in Washington. For. Ecol. Manage. 87, 233–256.

19. Dunlap, J.M., Heilman, P.E., and Stettler, R.F. (1994). Genetic variation and productivity of *Populus trichocarpa* and its hybrids. VII. Two-year survival and growth of native black cottonwood clones from four river valleys in Washington. Can. J. For. Res. 24, 1539–1549.

20. Newcombe, G., Stirling, B., McDonald, S., and Bradshaw, H.D. (2000). *Melampsora* × *columbiana*, a natural hybrid of *M. medusae* and *M. occidentalis*. Mycol. Res. 104, 261–274.

21. Barge, E.G., Leopold, D.R., Peay, K.G., Newcombe, G., and Busby, P.E. (2019). Differentiating spatial from environmental effects on foliar fungal communities of *Populus trichocarpa*. J. Biogeogr. 46, 2001–2011.

22. Lindow, S.E., and Brandl, M.T. (2003). Microbiology of the phyllosphere. Appl. Environ. Microbiol. 69, 1875–1883.

23. Vorholt, J.A. (2012). Microbial life in the phyllosphere. Nat. Rev. Microbiol. 10, 828–840.

24. Hacquard, S., Spaepen, S., Garrido-Oter, R., and Schulze-Lefert, P. (2017). Interplay between innate immunity and the plant microbiota. Annu. Rev. Phytopathol. 55, 565–589.

25. Jones, J.D.G., and Dangl, J.L. (2006). The plant immune system. Nature 444, 323–329.

26. Warton, D.I., Blanchet, F.G., O’Hara, R.B., Ovaskainen, O., Taskinen, S., Walker, S.C., and Hui, F.K.C. (2015). So many variables: joint modeling in community ecology. Trends Ecol. Evol. 30, 766–779.

27. Kainer, D., Hyatt, P.D., Jones, P., Streich, J., Shah, M., Ranjan, P., and Chen, J. (2020). The *Populus trichocarpa* pan-genome provides insights into the function of gene presence and absence variation. Nat. Genet. In press.

28. Zhang, B., Zhu, W., Diao, S., Wu, X., Lu, J., Ding, C., and Su, X. (2019). The Poplar pangenome provides insights into the evolutionary history of the genus. Commun. Biol. 2, 215.

29. Duplessis, S., Major, I., Martin, F., and Séguin, A. (2009). Poplar and pathogen interactions: insights from *Populus* genome-wide analyses of resistance and defense gene families and gene expression profiling. CRC. Crit. Rev. Plant Sci. 28, 309–334.

30. Wallace, J.G., Kremling, K.A., Kovar, L.L., and Buckler, E.S. (2018). Quantitative genetics of the maize leaf microbiome. Phytobiomes J. 2, 208–224.

31. Ke, P.-J., and Letten, A.D. (2018). Coexistence theory and the frequency-dependence of priority effects. Nat. Ecol. Evol. 2, 1691–1695.

32. Fukami, T., Mordecai, E.A., and Ostling, A. (2016). A framework for priority effects. J. Veg. Sci. 27, 655–657.

33. Peay, K.G., Belisle, M., and Fukami, T. (2012). Phylogenetic relatedness predicts priority effects in nectar yeast communities. Proc. R. Soc. B Biol. Sci. 279, 749–758.

34. Carbone, I., White, J.B., Miadlikowska, J., Arnold, A.E., Miller, M.A., Kauff, F., U’Ren, J.M., May, G., and Lutzoni, F. (2016). T-BAS: Tree-Based Alignment Selector toolkit for phylogenetic-based placement, alignment downloads and metadata visualization: an example with the Pezizomycotina tree of life. Bioinformatics 33, btw808.

35. Jumpponen, A., and Jones, K.L. (2009). Massively parallel 454 sequencing indicates hyperdiverse fungal communities in temperate *Quercus macrocarpa* phyllosphere. New Phytol. 184, 438–48.

36. Arnold, A.E., and Lutzoni, F. (2007). Diversity and host range of foliar fungal endophytes: are tropical leaves biodiversity hotspots? Ecology 88, 541–549.

37. Fukami, T. (2015). Historical contingency in community assembly: integrating niches, species pools, and priority effects. Annu. Rev. Ecol. Evol. Syst. 46, 1–23.

38. Johnson, N.C., Wilson, G.W.T., Bowker, M.A., Wilson, J.A., and Miller, R.M. (2010). Resource limitation is a driver of local adaptation in mycorrhizal symbioses. Proc. Natl. Acad. Sci. 107, 2093–2098.

39. Urban, M.C., and De Meester, L. (2009). Community monopolization: local adaptation enhances priority effects in an evolving metacommunity. Proc. R. Soc. B Biol. Sci. 276, 4129–38.

40. Morella, N.M., Weng, F.C.-H., Joubert, P.M., Metcalf, C.J.E., Lindow, S., and Koskella, B. (2020). Successive passaging of a plant-associated microbiome reveals robust habitat and host genotype-dependent selection. Proc. Natl. Acad. Sci. 117, 1148–1159.

41. Nemergut, D.R., Schmidt, S.K., Fukami, T., O’Neill, S.P., Bilinski, T.M., Stanish, L.F., Knelman, J.E., Darcy, J.L., Lynch, R.C., Wickey, P., et al. (2013). Patterns and processes of microbial community assembly. Microbiol. Mol. Biol. Rev. 77, 342–356.

42. Zamioudis, C., and Pieterse, C.M.J. (2012). Modulation of host immunity by beneficial microbes. Mol. Plant-Microbe Interact. 25, 139–150.

43. Van Wees, S.C., Van der Ent, S., and Pieterse, C.M. (2008). Plant immune responses triggered by beneficial microbes. Curr. Opin. Plant Biol. 11, 443–448.

44. Busby, P.E., Peay, K.G., and Newcombe, G. (2016). Common foliar fungi of *Populus trichocarpa* modify Melampsora rust disease severity. New Phytol. 209, 1681–1692.

45. Greischar, M. a, and Koskella, B. (2007). A synthesis of experimental work on parasite local adaptation. Ecol. Lett. 10, 418–34.

46. Gloor, G.B., Macklaim, J.M., Vu, M., and Fernandes, A.D. (2016). Compositional uncertainty should not be ignored in high-throughput sequencing data analysis. Austrian J. Stat. 45, 73.

47. Quinn, T.P., Erb, I., Richardson, M.F., and Crowley, T.M. (2018). Understanding sequencing data as compositions: an outlook and review. Bioinformatics 34, 2870–2878.

48. Guo, X., Zhang, X., Qin, Y., Liu, Y.-X., Zhang, J., Zhang, N., Wu, K., Qu, B., He, Z., Wang, X., et al. (2020). Host-associated quantitative abundance profiling reveals the microbial load variation of root microbiome. Plant Commun. 1, 100003.

49. Humphrey, P.T., and Whiteman, N.K. (2020). Insect herbivory reshapes a native leaf microbiome. Nat. Ecol. Evol. 4, 221–229.

50. Fukami, T. (2004). Assembly history interacts with ecosystem size to influence species diversity. Ecology 85, 3234–3242.

51. Orrock, J.L., and Fletcher Jr., R.J. (2005). Changes in community size affect the outcome of competition. Am. Nat. 166, 107–111.

52. Chase, J.M. (2010). Stochastic community assembly causes higher biodiversity in more productive environments. Science. 328, 1388–1391.

53. Kardol, P., Souza, L., and Classen, A.T. (2013). Resource availability mediates the importance of priority effects in plant community assembly and ecosystem function. Oikos 122, 84–94.

54. Leopold, D.R., Wilkie, J.P., Dickie, I.A., Allen, R.B., Buchanan, P.K., and Fukami, T. (2017). Priority effects are interactively regulated by top-down and bottom-up forces: evidence from wood decomposer communities. Ecol. Lett. 20, 1054–1063.

55. Bodenhausen, N., Bortfeld-Miller, M., Ackermann, M., and Vorholt, J.A. (2014). A synthetic community approach reveals plant genotypes affecting the phyllosphere microbiota. PLoS Genet. 10, e1004283.

56. Vorholt, J.A., Vogel, C., Carlström, C.I., and Müller, D.B. (2017). Establishing causality: opportunities of synthetic communities for plant microbiome research. Cell Host Microbe 22, 142–155.

57. Christian, N., Whitaker, B.K., and Clay, K. (2015). Microbiomes: unifying animal and plant systems through the lens of community ecology theory. Front. Microbiol. 6, 1–15.

58. Nagelkerke, N.J.D. (1991). A note on a general definition of the coefficient of determination. Biometrika 78, 691–692.

59. Huang, Y.-L., Zimmerman, N., and Arnold, A. (2018). Observations on the early establishment of foliar endophytic fungi in leaf discs and living leaves of a model woody angiosperm, *Populus trichocarpa* (Salicaceae). J. Fungi 4, 58.

60. Toju, H., Tanabe, A.S., and Ishii, H.S. (2016). Ericaceous plant-fungus network in a harsh alpine-subalpine environment. Mol. Ecol. 25, 3242–3257.

61. Gardes, M., and Bruns, T.D. (1993). ITS primers with enhanced specificity for basidiomycetes - application to the identification of mycorrhizae and rusts. Mol. Ecol. 2, 113–8.

62. Ihrmark, K., Bödeker, I.T.M., Cruz-Martinez, K., Friberg, H., Kubartova, A., Schenck, J., Strid, Y., Stenlid, J., Brandström-Durling, M., Clemmensen, K.E., et al. (2012). New primers to amplify the fungal ITS2 region–evaluation by 454-sequencing of artificial and natural communities. FEMS Microbiol. Ecol. 82, 666–77.

63. Lundberg, D.S., Yourstone, S., Mieczkowski, P., Jones, C.D., and Dangl, J.L. (2013). Practical innovations for high-throughput amplicon sequencing. Nat. Methods 10, 999–1002.

64. Stevens, J.L., Jackson, R.L., and Olson, J.B. (2013). Slowing PCR ramp speed reduces chimera formation from environmental samples. J. Microbiol. Methods 93, 203–205.

65. Hamady, M., Walker, J.J., Harris, J.K., Gold, N.J., and Knight, R. (2008). Error-correcting barcoded primers for pyrosequencing hundreds of samples in multiplex. Nat. Methods 5, 235–237.

66. Somervuo, P., Koskinen, P., Mei, P., Holm, L., Auvinen, P., and Paulin, L. (2018). BARCOSEL: a tool for selecting an optimal barcode set for high-throughput sequencing. BMC Bioinformatics 19, 257.

67. Galanti, L., Shasha, D., and Gunsalus, K. (2017). Pheniqs: fast and flexible quality-aware sequence demultiplexing. bioRxiv, 128512.

68. Martin, M. (2011). Cutadapt removes adapter sequences from high-throughput sequencing reads. EMBnet.journal 17, 10.

69. Sturm, M., Schroeder, C., and Bauer, P. (2016). SeqPurge: highly-sensitive adapter trimming for paired-end NGS data. BMC Bioinformatics 17, 208.

70. Callahan, B.J., McMurdie, P.J., Rosen, M.J., Han, A.W., Johnson, A.J.A., and Holmes, S.P. (2016). DADA2: High-resolution sample inference from Illumina amplicon data. Nat. Methods 13, 581–583.

71. Wright, Erik S. (2016). Using DECIPHER v2.0 to analyze big biological sequence data in R. R J. 8, 352.

72. Tuskan, G.A., DiFazio, S., Jansson, S., Bohlmann, J., Grigoriev, I., Hellsten, U., Putnam, N., Ralph, S., Rombauts, S., Salamov, A., et al. (2006). The genome of Black Cottonwood, *Populus trichocarpa* (Torr. & Gray). Science. 313, 1596–1604.

73. Langmead, B., and Salzberg, S.L. (2012). Fast gapped-read alignment with Bowtie 2. Nat. Methods 9, 357–359.

74. McMurdie, P.J., and Holmes, S. (2013). phyloseq: An R package for reproducible interactive analysis and graphics of microbiome census data. PLoS One 8, e61217.

75. Untergasser, A., Cutcutache, I., Koressaar, T., Ye, J., Faircloth, B.C., Remm, M., and Rozen, S.G. (2012). Primer3—new capabilities and interfaces. Nucleic Acids Res. 40, e115–e115.

76. Lofgren, L.A., Uehling, J.K., Branco, S., Bruns, T.D., Martin, F., and Kennedy, P.G. (2019). Genome-based estimates of fungal rDNA copy number variation across phylogenetic scales and ecological lifestyles. Mol. Ecol. 28, 721–730.

77. McLaren, M.R., Willis, A.D., and Callahan, B.J. (2019). Consistent and correctable bias in metagenomic sequencing experiments. Elife 8, 559831.

78. Schindelin, J., Arganda-Carreras, I., Frise, E., Kaynig, V., Longair, M., Pietzsch, T., Preibisch, S., Rueden, C., Saalfeld, S., Schmid, B., et al. (2012). Fiji: an open-source platform for biological-image analysis. Nat. Methods 9, 676–682.

79. Arganda-Carreras, I., Kaynig, V., Rueden, C., Eliceiri, K.W., Schindelin, J., Cardona, A., and Sebastian Seung, H. (2017). Trainable Weka Segmentation: a machine learning tool for microscopy pixel classification. Bioinformatics 33, 2424–2426.

80. Wang, Y., Naumann, U., Wright, S., and Warton, D. (2019). mvabund: statistical methods for analysing multivariate abundance data. R package version 4.0.1. URL https://CRAN.R-project.org/package=mvabund.

81. Warton, D.I. (2011). Regularized sandwich estimators for analysis of high-dimensional data using generalized estimating equations. Biometrics 67, 116–123.

82. Warton, D.I., Thibaut, L., and Wang, Y.A. (2017). The PIT-trap—A “model-free” bootstrap procedure for inference about regression models with discrete, multivariate responses. PLoS One 12, e0181790.

83. Venables, W.N., and Ripley, B.D. (2002). Modern Applied Statistics with S Fourth Edi. (Springer New York).

84. Vannette, R.L., and Fukami, T. (2014). Historical contingency in species interactions: towards niche-based predictions. Ecol. Lett. 17, 115–124.

85. DiCiccio, T., and Efron, B. (1996). Bootstrap confidence intervals. Stat. Sci. 11, 189–212.

86. Hacquard, S., Petre, B., Frey, P., Hecker, A., Rouhier, N., and Duplessis, S. (2011). The Poplar-Poplar rust interaction: insights from genomics and transcriptomics. J. Pathog. 2011, 1–11.

87. Cribari-Neto, F., and Zeileis, A. (2010). Beta Regression in R. J. Stat. Softw. 34, 1–24.

88. Smithson, M., and Verkuilen, J. (2006). A better lemon squeezer? Maximum-likelihood regression with beta-distributed dependent variables. Psychol. Methods 11, 54–71.

89. Zeileis, A., and Hothorn, T. (2002). Diagnostic checking in regression relationships. R News 2, 7–10.

90. R Core Team (2019). R: A language and environment for statistical computing. R Foundation for Statistical Computing, Vienna, Austria. URL https://www.R-project.org/.

