## Supplemental Materials for "Host genotype and colonist arrival order jointly govern plant microbiome composition and function"

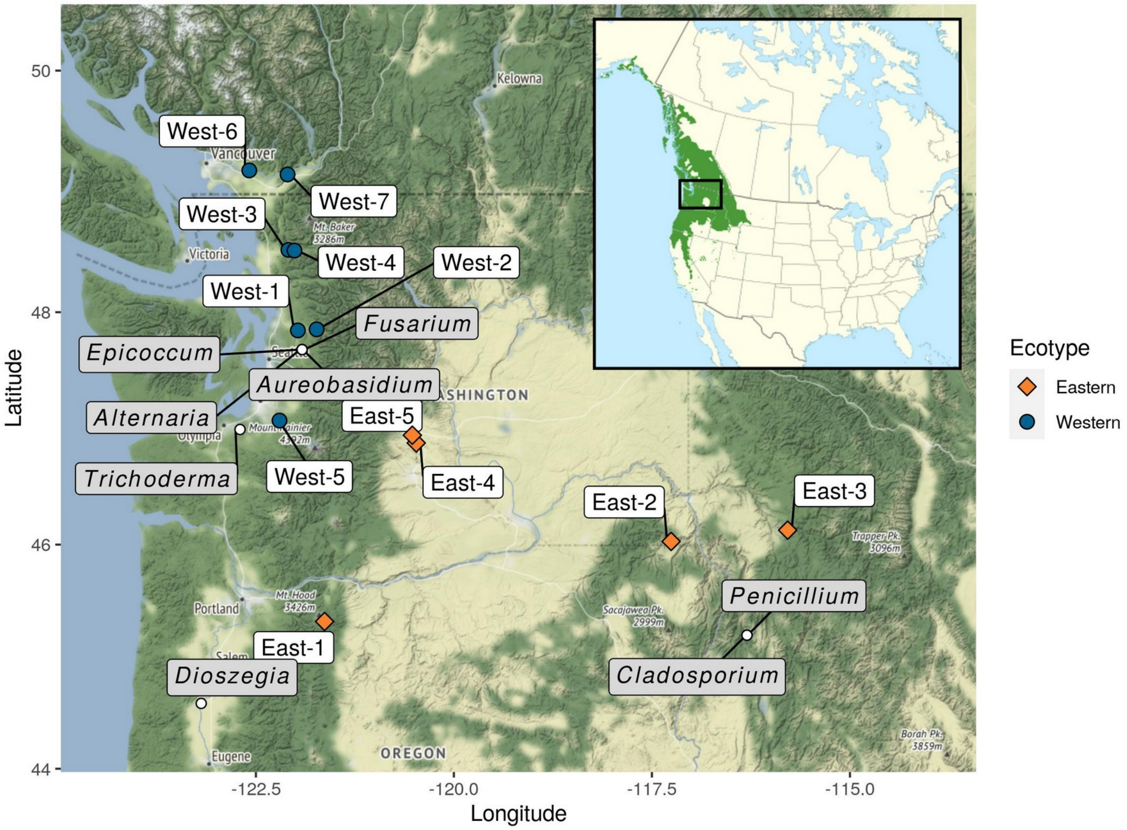


**Figure S1: Origins of *P. trichocarpa* genotypes and fungal isolates used in the greenhouse experiment. Related to Star Methods.**

*P. trichocarpa* genotypes used in the greenhouse experiment were originally collected from populations located both east and west of the Cascade Divide, within the core of the species' geographic range in northwest North America (inset). Fungal isolates were isolated from *P. trichocarpa* leaves collected from multiple locations.

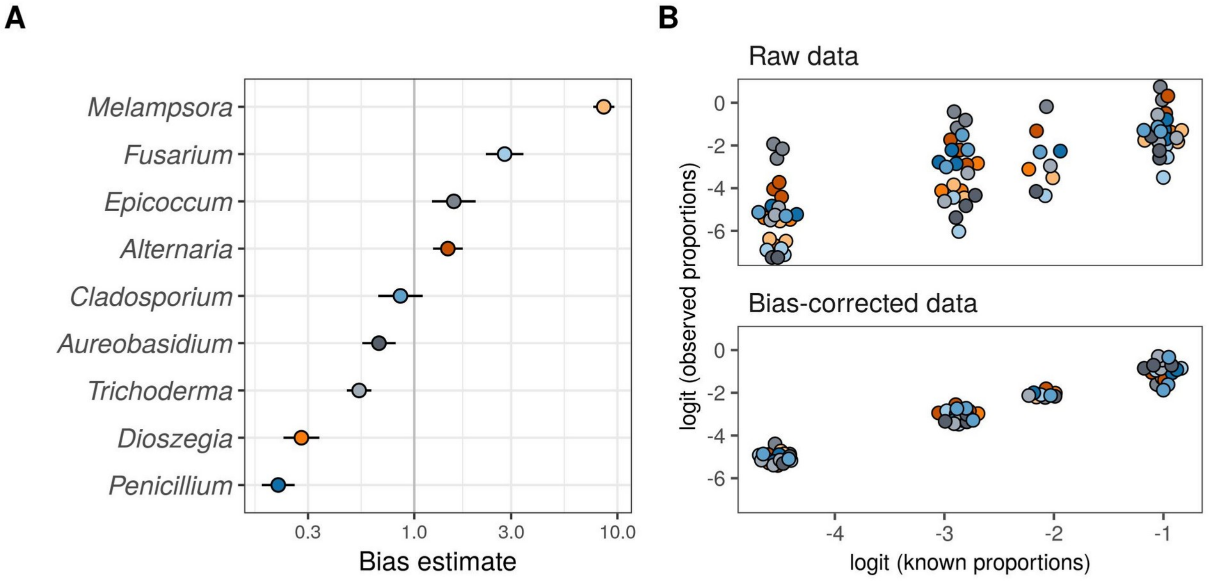


**Figure S2: Estimates of taxon specific sequencing bias used in analyses of fungal community composition. Related to Star Methods.**

(A) Point estimates of species-specific sequencing bias and bootstrap estimated confidence intervals (+/- two geometric standard errors) calculate from 10 mock community samples. Larger values indicate greater observed relative abundance than expected.

(B) The relationship between the observed proportional abundance of sequence reads and the known proportional abundance of template DNA (genome copy number) in the 10 mock community samples before (top) and after (bottom) applying bias-correction. Each point represents the proportional abundance of one species (indicated by point colors) in one mock community sample.

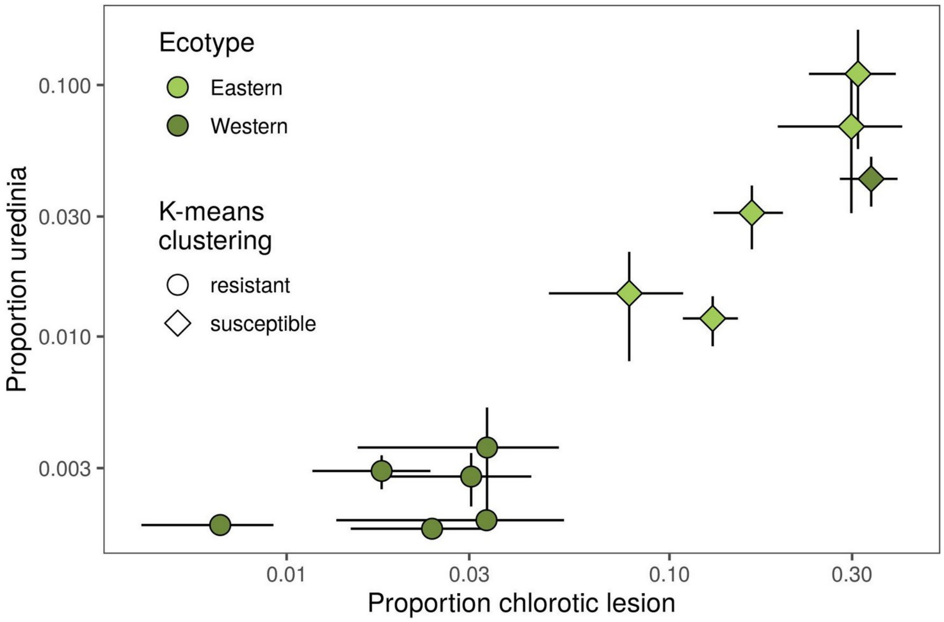


**Figure S3: Hierarchical clustering of *P. trichocarpa* rust disease susceptibility to identify susceptible genotypes for rust severity analyses. Related to Figure 4.**
Control *P. trichocarpa* plants, inoculated with only the *Melampsora* rust pathogen, were used to identify rust resistant (cluster 1, circles) and rust susceptible (cluster 2, diamonds) plant genotypes. Here, the results of K-means clustering of $log_{10}$-transformed measures of disease severity (proportion of leaf area occupied by chlorotic lesions or rust uredinia) are shown, confirming our expectation that western ecotypes are often, but not always, more resistant to *Melampsora* leaf rust than eastern ecotypes. Points show mean rust severity (± se), weighted to account for variation in the total leaf area sampled for each of 5 clonal replicates.

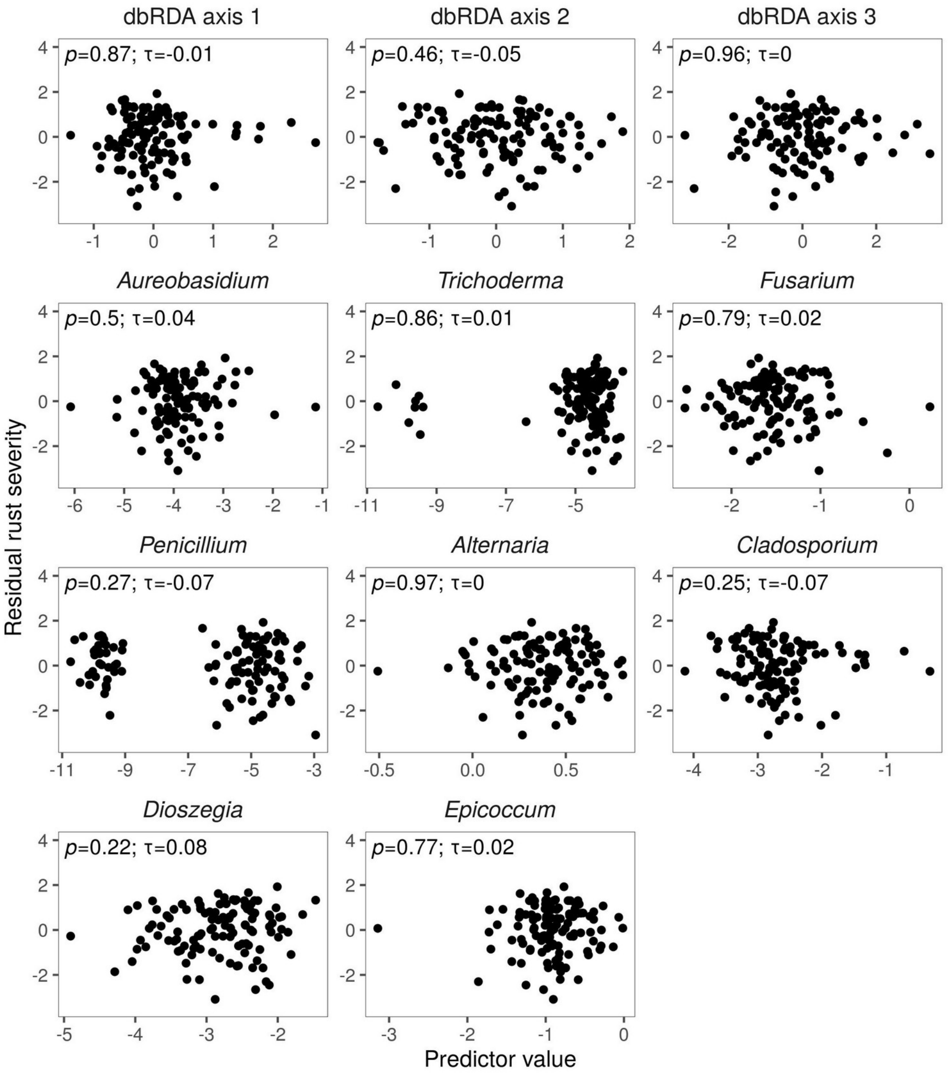


**Figure S4: Rust disease severity is not correlated with fungal species composition or the relative abundance of individual taxa. Related to Figure 4.**

Residual variation in rust disease severity, after accounting for variation among plant genotypes (y-axis), is not correlated with variation in fungal community composition associated with fungal species arrival order (top, dbRDA axes 1-3) or, variation in the proportional abundance of individual foliar fungi. Statistics presented in each panel show the results of a correlation test using Kendall's tau.

| | Genotype | | | | Ecotype | | | |
|---|---|---|---|---|---|---|---|---|
| | Df | Df.resid | Wald-$\chi^2$ | *P*-value | Df | Df.resid | Wald-$\chi^2$ | *P*-value |
| Host | 11 | 235 | 20.2 | < 0.001 | 1 | 245 | 12.4 | < 0.001 |
| Treatment | 4 | 231 | 21.8 | < 0.001 | 4 | 241 | 21.7 | < 0.001 |
| Host:Treatment | 44 | 187 | 23.5 | 0.038 | 4 | 237 | 6.2 | 0.302 |

**Table S1: Full results of multivariate, negative-binomial generalized linear models of fungal community composition. Related to Figure 2.**

Models tested whether fungal species composition in the *P. trichocarpa* foliar microbiome varied with species arrival order treatments, and whether these effects vary among individual plant genotypes, or ecotypes (i.e., originating from east or west of the Cascade Divide). *P*-values are based on permutation tests and Wald test statistics accounting for possible correlations among taxa.

| | Susceptible | | | Resistant | | |
|---|---|---|---|---|---|---|
| | ΔDf | LRT-$\chi^2$ | *P*-value | ΔDf | LRT-$\chi^2$ | *P*-value |
| Genotype | 5 | 60.2 | < 0.001 | 5 | 33.5 | 0.000 |
| Treatment | 4 | 4.5 | 0.339 | 4 | 1.7 | 0.784 |
| Genotype:Treatment | 20 | 33.6 | 0.029 | 20 | 5.9 | 0.999 |

**Table S2: Full model results of beta-regression models of rust severity. Related to Figure 4.** Likelihood ratio tests comparing beta-regression models of rust severity for both rust susceptible and rust resistant *P. trichocarpa* genotypes.

| Isolate ID | Genus ID | Genbank accession # | Nearest UNITE Species Hypothesis | Notes |
|---|---|---|---|---|
| PE_29 | *Alternaria* | MT035960 | *Alternaria alternata* (Fr.) Keissl. \| SH2179670.08FU | |
| PE_11 | *Aureobasidium* | MT035961 | *Aureobasidium pullulans* (de Bary) G. Arnaud \| SH2149988.08FU | |
| PE_07 | *Cladosporium* | MT035962 | *Cladosporium* Link \| SH2320293.08FU (99) | The only mismatch with the reference sequence in an ambiguous base (N) in the database sequence; effectively a 100% match. |
| Y_23 | *Dioszegia* | MT035963 | *Dioszegia butyracea* Q.M. Wang & F.Y. Bai \| SH2454824.08FU | |
| 12_41 | *Epicoccum* | MT035964 | Pezizomycotina \| SH2232033.08FU | Chlamydospore morphology suggests that the correct species is *Epicoccum nigrum*. This is also the taxonomic identification given to the representative sequence of this Species Hypothesis. |
| PE_12 | *Fusarium* | MT035965 | *Gibberella pulicaris* (Fr.) Sacc. \| SH2230102.08FU<br>Nectriaceae \| SH2229701.08FU | Only the ITS1 region was available due to failed Sanger sequencing, resulting in multiple 100% matches in the UNITE database. Most matching Species Hypotheses are dominated by plant endophytes and pathogens often identified as *Fusarium avenaceum* or *Fusarium tricinctum*. Chlamydospore morphology confirms *Fusarium* sp. |
| 2_85 | *Penicillium* | MT035966 | *Penicillium bialowiezense* K.M. Zalessky \| SH2189921.08FU | |
| 12-65 | *Trichoderma* | MT035967 | *Trichoderma trixiae* Samuels & Jaklitsch \| SH2303501.08FU | |

**Table S3: Identifying information and putative taxonomy of the fungal isolates used in the current experiment. Related to Star Methods.**

Taxonomic predictions were made by manual curation against the UNITE Species Hypothesis database (v8.2).
